# Pathological Angiogenesis Precedes the Onset of Aortic Dissection

**DOI:** 10.1101/2025.09.10.675317

**Authors:** Kazuaki Yoshioka, Kenji Iino, Yukinobu Ito, Tomohiro Iba, Takahisa Matsuzaki, Nanami Sakurai, Masafumi Horie, Tomoaki Matsuyama, Taku Wakabayashi, Aya Matsui, Yoshitaka Yamamoto, Takashi Satoh, Akiteru Goto, Hiroshi Y. Yoshikawa, Hitoshi Ando, Daichi Maeda, Hirofumi Takemura, Hisamichi Naito

## Abstract

Aortic dissection (AD) is a severe, life-threatening disease that occurs abruptly^1^. Although remodeling and inherent vulnerability of the aortic media, as observed in heritable connective tissue disorders such as Marfan syndrome, have been implicated in the onset of AD, the precise pathological mechanisms underlying the non-heritable form, which accounts for most cases, remains largely unknown^2^. Here, using hyperacute AD patient-derived samples, we show that pathological angiogenesis within the aortic media precedes the onset of dissection. Integrating single-cell transcriptomic analysis of aortic medial endothelial cells with high-resolution imaging revealed temporal changes preceding dissection. Hypoxic alterations in the media triggered osteochondrogenic changes in smooth muscle cells, promoted hydroxyapatite deposition, and induced angiogenesis via VEGF secretion from AD-specific CD14^+^CD68^+^CD163^+^MRC1(CD206)^neg-low^ macrophages. Atomic force microscopy revealed that these processes culminated in the formation of a soft capillary layer, potentially concentrating stress on this region and contributing to AD onset. H&E staining of archived specimens revealed that non-heritable AD can be classified into angiogenic or non-angiogenic subtypes, with more than half exhibiting the former. In contrast, acute AD cases associated with Marfan syndrome typically exhibit a non-angiogenic pattern, suggesting distinct underlying mechanisms. This study reports angiogenesis as a key pathological driver in non-heritable AD and highlight potential targets for early diagnosis, prevention, and treatment.

## Introduction

Aortic dissection (AD) is a sudden-onset, life-threatening condition first described over 250 years ago but remaining a major cause of clinical emergencies today ^1,3^. Approximately 40% of patients die before reaching the hospital; if the ascending aorta is involved, the mortality rate without surgical intervention exceeds 80% within two weeks ^4^. AD results from a separation of the aortic wall within the media, typically between the inner two-thirds and outer one-third, and usually involves the ascending aorta. Medial degeneration, which results from matrix degradation and inflammation, is widely considered to be a major factor weakening the aortic wall ^2^. While heritable connective tissue disorders such as Marfan syndrome (fibrillin-1 mutation) and Loeys-Dietz syndrome (TGFβ signaling defects) are established contributors to medial degeneration ^5,6^, they account for only about 5% of AD cases ^7,8^. In contrast, in most non-heritable AD, the mechanisms of medial degeneration remain largely unknown. Dissection may be initiated by an intimal tear or by intramural hematoma resulting from rupture of the vasa vasorum ^9,10^, or is occasionally associated with trauma, leading to tearing of the media. However, these events likely represent only a part of a more complex pathological process.

Angiogenesis, the formation of new blood vessels from pre-existing ones, is a vital process during development and a hallmark of many diseases including cancer, atherosclerosis and diabetic retinopathy ^11^. Under physiological conditions it occurs infrequently in adults, except in the female reproductive tract. Angiogenesis is typically induced by hypoxia triggering the secretion of vascular endothelial growth factor (VEGF), which activates endothelial cells (ECs) through VEGF receptors (VEGFRs), and leads to the emergence of tip cells and proliferating stalk cells ^12^.

These angiogenic ECs have been characterized histologically and by single-cell RNA sequencing (scRNA-seq) ^13,14^. Macrophages are known stimulators of angiogenesis ^15^. In several different models, new vessel formation in adults requires more than two days to occur ^16–18^.

Aortic aneurysm represents another major type of aortic disease. A typical true aortic aneurysm is chronic aortic disease defined as a localized dilatation of the aorta with a diameter at least 50% greater than normal, and is contained by all three layers of the aortic wall ^19^. Multiple pathological processes, including inflammation and immune responses, degradation of aortic wall connective tissue, phenotype switching of vascular smooth muscle cells (SMCs), oxidative stress, biomechanical wall stress and neovascularization, contribute to this process ^20^. In contrast, only minimal structural abnormalities have traditionally been observed in the aortic wall prior to the onset of non-heritable AD ^21^.

Here, by integrating temporal analyses of cellular dynamics and protein expression with data from 22 freshly obtained hyperacute AD samples (collected within 12 hours post-onset) and 34 archived hyperacute cases, we demonstrate that pathological angiogenesis of the vasa vasorum is a key mechanism underlying the onset of many non-heritable aortic dissections. We identified a marked increase in medial blood vessels, with ECs exhibiting tip-like and proliferative phenotypes.

CD163⁺ macrophages were found to express VEGF, acting on ECs through VEGFRs. We also found medial hypoxia, osteochondrogenic changes in SMCs, and hydroxyapatite deposition. Pathological blood vessels, along with these associated changes, appear to weaken the medial wall and create regions that are particularly susceptible to tearing. Notably, these angiogenic changes were commonly but not always observed in non-heritable AD, but less often in Marfan syndrome-associated cases. Our findings indicate that non-heritable AD can be subclassified into angiogenic and non-angiogenic types, with more than 50% of cases exhibiting the former, suggesting that pathological angiogenesis in the media is a critical event that precedes the onset of many ADs. These findings help reshape the current understanding of AD and may guide future diagnostic and therapeutic strategies.

## Results

### Pathological Angiogenesis in the Aortic Wall

To investigate the molecular mechanism underlying AD, we obtained fresh dissected ascending aorta from non-heritable hyperacute AD patient who underwent surgical artificial blood vessel replacement within 12 hours after the onset (data not shown). We performed scRNA-seq analysis on CD31^+^CD45^-^ cells—ECs of vasa vasorum—isolated from the outer layer of dissected aorta, which includes the media and adventitia (data not shown). Dimensionality reduction by UMAP identified 16 clusters (amongst 6,028 cells), of which 10 contained vascular EC markers (positive for *CDH5*, *PECAM1*, *EMCN* and *ERG*, but negative for *PTPRC* and *PROX1*). To validate cluster identity of vascular ECs, we used canonical marker genes and the top 10 differentially expressed genes (DEGs) to confirmed 3 venous ECs (cluster 0, 1 and 2 marked by *ACKR1*, *NR2F2* and NRP2), post-capillary venule-like ECs (cluster 5 marked by *CPE* and *POSTN*), capillary ECs (cluster 4 marked by *RGCC* and *SPARC*), arterial ECs (cluster 6 marked by *GJA4*, *GJA5*, *BMX*, *FBLN5*, *SOX17* and *HEY1*), tip-like ECs (cluster 7 marked by *ESM1*, *APLN*, *DLL4* and *CXCR4*), immature ECs (cluster 3 marked by *APLNR* and *IGFBP5*), proliferating ECs (cluster 8 marked by *MKI67* and *PCLAF*), and *ERG*^high^-undefined ECs (cluster 10 marked by *ERG*) ^14,22^ (Fig. 1a-c, and data not shown). Compared to non-dissected control aorta obtained from distal arch aneurysm patient (7,999 cells) (Fig. 1d), we found striking differences in cell distribution patterns in the tip-like, proliferating, and immature ECs clusters, suggesting a predominant angiogenic gene expression pattern in ECs from the AD sample (Fig. 1e-g, and data not shown).

**Figure 1.**
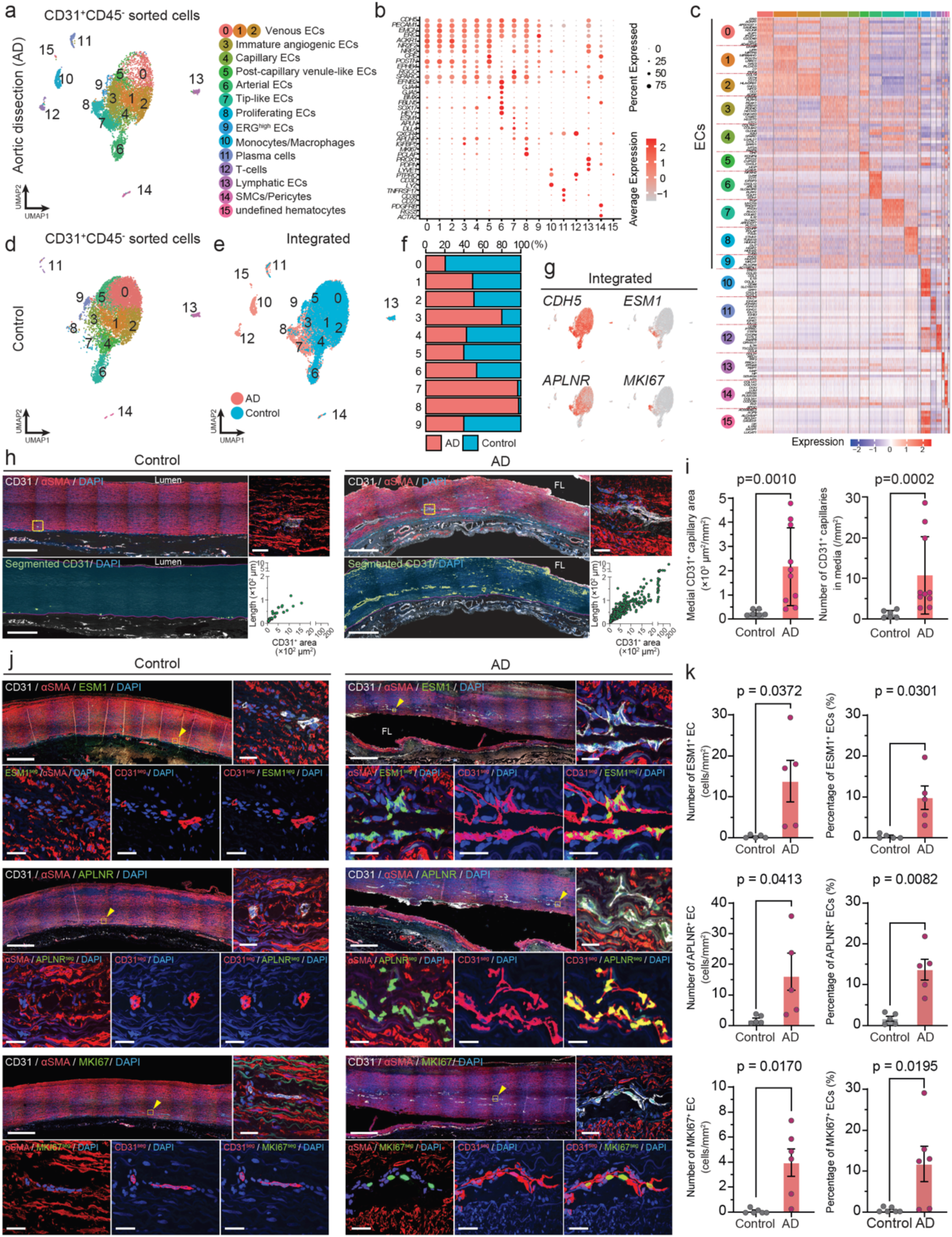
Pathological Angiogenesis of the Vasa Vasorum in the Aortic Media of Aortic Dissection. **(a)** UMAP plots of scRNA-seq data from 6,028 CD31^+^CD45^-^ sorted cells isolated from aortic media and adventitia of an AD sample. (**b**) Bubble plots comparing the expression levels of selected signature genes across clusters in the CD31^+^CD45^-^ sorted cells from the AD sample. (**c**) Heatmap showing the expression levels of the top ten differentially expressed genes (DEGs) in each cluster of CD31^+^CD45^-^ sorted cells from the AD sample. (**d**) UMAP plots of scRNA-seq data from CD31⁺CD45⁻ sorted cells isolated from the aortic media and adventitia of a control sample. (**e**) Integrated UMAP plots of CD31⁺CD45⁻ sorted cells from AD and control samples, color-coded by sample origin. (**f**) Relative proportions of AD and control within each EC cluster. (**g**) Feature plots of selected genes projected onto the integrated UMAP plots of CD31⁺CD45⁻ sorted cells. (**h**) Representative immunofluorescence images of aortic sections stained for CD31 (white) and αSMA (red), with magnified views of the yellow-boxed regions. Nuclei were counterstained with DAPI (blue). Segmented images of CD31-positive capillaries (green) are shown in the lower panels. Scatter plots display pixel-based area and length of the segmented CD31-positive capillaries. TL, true lumen; FL, false lumen; the dotted line indicates the border between the media and adventitia. Scale bars: 1 mm in wide-field images and 20 µm in magnified images. (**i**) Bar plots show the quantification results for the media within the purple-lined ROI. n = 5–8; data are expressed as mean ± S.D. (**j**) Representative immunofluorescence images of aortic sections stained for CD31 (white), αSMA (red), and ESM1 (green, top), APLNR (green, middle), or MKI67 (green, bottom), with magnification of the yellow-boxed regions depicted by arrowheads. Nuclei were counterstained with DAPI (blue). Segmented images of CD31 (CD31^seg^, red), ESM1 (ESM1^seg^, green), APLNR (APLNR^seg^, green), or MKI67 (MKI67^seg^, green), and double-positive cells (yellow) are shown in the lower panels. Scale bars: 1 mm in wide-field images and 20 µm in magnified images. (**k**) Bar plots show the quantification results of the number (left) and percentage (right) of ESM1^+^, APLNR^+^, and MKI67^+^-endothelial cells (ECs) in the media. n = 6–10; data are expressed as mean ± S.D.

We validated the angiogenic characteristics in AD using orthogonal approaches. First, we evaluated vasa vasorum in the media and found that relative to controls, there were significantly greater numbers and areas of CD31^+^ blood vessels with robust sprouting in the outer layer of media in AD samples obtained within 12 hours after onset (Fig.1h, i, and data not shown). This was unexpected, as the formation of new blood vessels after a pathological event typically takes more than 2 days ^23^. Indeed, a rodent hind-limb ischemia model demonstrated that the increase in blood vessels occurs 2 to 3 days after the induction of ischemia (data not shown). Moreover, to confirm that the increase in vessel number was not merely due to the dilation of small capillaries or closed arteriovenous shunts, we quantified ECs by counting their nuclei and found a significant increase in their numbers in AD specimens (data not shown). Importantly, the increase in the number of blood vessels in the media was greatest in the outer one-third, around the site of dissection, i.e., the false lumen, with some parts of the false lumen even positive for CD31. We next performed immunostaining for ESM1, APLNR and MKI67 to confirm the presence of tip-like, immature and proliferating ECs and observed a significant and specific increase in these cells, suggestive of pathological angiogenesis (Fig. 1j, k). We further found increases of blood vessels in the adventitia and surrounding perivascular adipose tissue in AD samples (data not shown). It is important to note that not all individual cases exhibited such vascular proliferation. Although many samples (the proportion of which will be discussed in detail later) showed increased vascularity, some displayed minimal expression of ESM1, MKI67 and APLNR by immunostaining (data not shown). In the latter, a small number of tip cells was detected without forming a clear cluster, and proliferating cells were not observed, while an *IGFBP5*-expressing immature cluster was detectable in the single-cell analysis of ECs (data not shown). Taken together, these findings suggest that pathological angiogenesis in the outer layer of the media precedes the onset of dissection in many cases. We also identified two patterns of pathological angiogenesis: one characterized by increased vascularity accompanied by active angiogenesis, and the other with increased vascularity but low angiogenic activity.

### Macrophages induce angiogenesis

Next, focusing on the cells surrounding ECs, we explored the mechanisms whereby angiogenesis is induced. We simultaneously isolated whole cells from the lysate of dissected outer layer, which was used for scRNA-seq analysis of ECs, and independently performed scRNA-seq analysis. Unbiased clustering of 4503 cells revealed 15 cell clusters including 3 EC clusters (cluster 0,1 and 2 marked by *PECAM1*, *CDH5*), pericyte/SMCs (cluster 3 marked by *MYH11*, *ACTA2*, *PDGFRB* and *RGS5*), 3 fibroblast clusters (cluster 4, 5 and 6 marked by *COL1A1*, *COL1A2*, *FBLN1*, *PDGFRA*), and 8 immune cell lineages, including 3 monocyte/macrophage populations (cluster 7,8 and 9 marked by *LYZ*, *ITGAM, CD68*), T-lymphocytes (cluster 10 marked by *CD3D*), natural killer cells (cluster 11 marked by *NKG7*), B-lymphocytes (cluster 12 marked by *CD27* and *MS4A1*), plasma cells (cluster 13 marked by *TNFRSF17* and *JCHAIN*), and neutrophils (cluster 14 marked by *S100A8/9* and *FCGR3B*) (Fig. 2a and data not shown). Compared to the non-dissected control aorta, we noted the appearance of a neutrophil cluster and an increase in the proportion of macrophages, suggesting the migration of inflammatory cells into the vascular wall (Fig. 2b-e, and data not shown).

**Figure 2.**
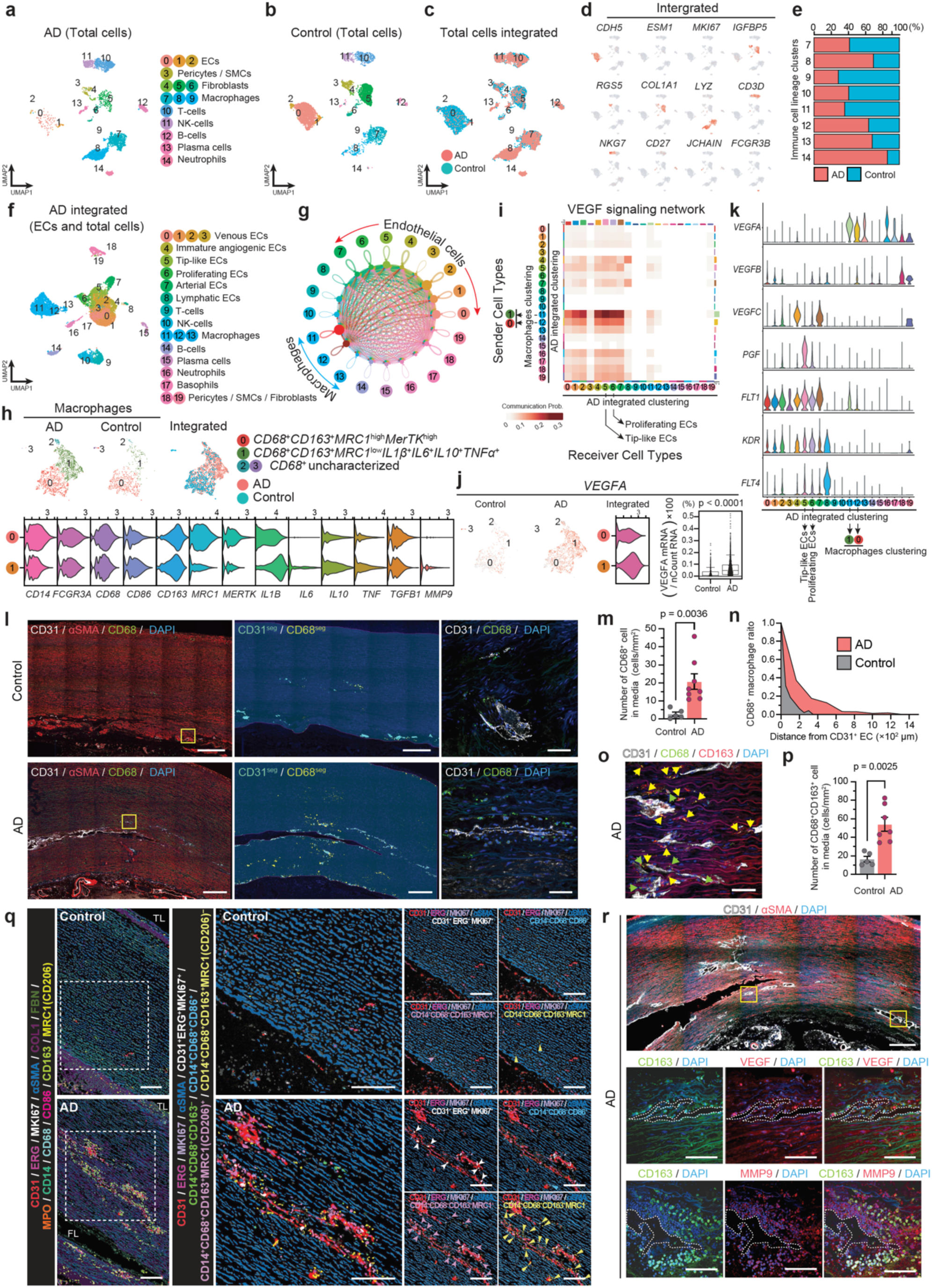
AD-Associated Macrophages with High Cytokine Secretion Induce Medial Angiogenesis via VEGFA. (**a** to **c**) UMAP plots of scRNA-seq data from total cells of aortic media and adventitia from AD (4,503 cells) (**a**) and control (5,754 cells) (**b**). Integrated UMAP plots of total cells from both AD and control samples, color-coded by sample origin (**c**). (**d**) Feature plots of the indicated genes projected onto the integrated UMAP plots of total cells. (**e**) Relative proportions of AD and control within each immune cell lineage cluster. (**f**) Integrated UMAP plots of CD31⁺CD45⁻ sorted cells and total cells from AD patients. (**g**) Cell–cell communications analysis between each cluster by CellChat analysis based on interaction strength and numbers. (**h**) UMAP plots of macrophages extracted from total cells of AD and control samples, and their integrated dataset. Violin plots showing the expression of selected macrophage markers and cytokines in subclusters 0 and 1 from the integrated data of extracted macrophages. (**i**) Heatmap of the VEGF signaling network between the two cell types. (**j**) Feature plots of VEGFA expression in macrophages of control and AD samples, and violin plots of VEGFA expression in subclusters 0 and 1 from the integrated data of extracted macrophages. Graphs show the distribution of the percentage of VEGFA gene transcripts per cell. (**k**) Violin plots show the expression of selected genes representing VEGF signaling across each cluster. (**l**) Representative immunofluorescence images of aortic sections stained for CD31 (white), αSMA (red), and CD68 (green), counterstained with DAPI (blue). Segmented images of CD31-positive blood vessels (blue) and CD68-positive macrophages (yellow) are shown in the middle panels. Magnified views of the yellow-boxed regions in the left panels are shown in the right panels. Scale bars: 500 µm in wide-field images and 50 µm in magnified images. (**m**) Bar plot showing quantification of CD68-positive macrophages in the media. n = 5–8; data are expressed as mean ± S.D. (**n**) Representative histogram of the distribution of macrophage-to-endothelial cell distance in AD and control. (**o**) Representative immunofluorescence image of dissected aortic sections stained for CD31 (white), CD68 (green), and CD163 (red), counterstained with DAPI (blue). Green arrows: CD68 single-positive cells; yellow arrows: CD68 and CD163 double-positive cells. Scale bars: 50 µm. (**p**) Bar plot showing quantification of CD68 and CD163 double-positive macrophages in the media. n = 5–7; data are expressed as mean ± S.D. (**q**) Representative images from imaging mass cytometry (IMC) analysis of aortic sections stained for CD31, ERG, MKI67, αSMA, Collagen Type I, Fibronectin, MPO, CD14, CD86, CD163, and MRC1 (CD206). Images of each segmented marker are shown in the left panels. The middle and right panels display magnified views of the dashed boxes in the left panels. Segmented images of MKI67-positive proliferating ECs and specific types of macrophages are also shown. Scale bars: 100 µm in wide-field images and 50 µm in magnified images. (**r**) The upper panel shows a representative immunofluorescence image of dissected aortic sections stained for CD31 (white) and αSMA (red), counterstained with DAPI (blue). Magnified views of the regions indicated by the yellow boxes in the upper panel are shown in the middle and lower panels. The middle panels are stained for CD163 (green) and VEGF (red), while the lower panels are stained for CD163 (green) and MMP9 (red), both counterstained with DAPI (blue). Scale bars: 500 µm in the upper panel and 100 µm in the middle and lower panels.

Next, we investigated the interactions between each EC cluster and the surrounding cells based on ligand-receptor expression patterns ^24,25^. To achieve this, we integrated the whole-cell scRNA-seq dataset with the EC scRNA-seq dataset (11,515 cells) (Fig. 2f and data not shown). This enabled us to precisely analyze cell-cell interactions between the cells of each EC cluster and their surrounding cells.

Comprehensive analysis with whole-cell clusters showed a large number of signals between the macrophage clusters and the ECs clusters including the tip-like cells, proliferating ECs, and immature angiogenic ECs (Fig 2g and data not shown). We therefore focused on macrophages, extracted them from the scRNAseq data, and compared them between AD (2,163 cells) and control (721 cells). In the latter, resident macrophages were characterized as *CD14*⁺*CD68*⁺*CD80*^-^ *CD163*⁺*MRC1(CD206*)^high^*MerTK*^high^ (corresponding to cluster 12 in Fig. 2f), whereas in AD, together with their increase, we identified a distinct AD-specific macrophage cluster—*CD68*^+^*CD163*^+^*MRC1(CD206)*^neg-low^*MerTK*^low^—which expresses the pro-inflammatory cytokines *IL1β*, *IL6*, and *TNFα*, as well as the anti-inflammatory cytokine *IL10* (Fig. 2h and data not shown). Focusing on VEGF signaling between macrophages and EC clusters, we found that *VEGFA* was more abundantly expressed in the AD-specific macrophage cluster and acted as a ligand for EC clusters via *VEGFR1* and *VEGFR2* (Fig. 2i-k). We further found that these macrophages exhibit increased expression of *MMP9* and *MMP19* (data not shown), suggesting an active remodeling role during angiogenesis ^26^.

We next validated the cell-cell interactions by immunostaining. In the media of AD samples, we found that CD68⁺ macrophages were significantly increased compared to controls (Fig. 2l, m). Notably, we observed an increase in macrophages not only around the pathological medial vessels in AD samples, but also in areas distant from these vessels—a phenomenon rarely observed in controls (Fig. 2n).

Furthermore, most CD68⁺ macrophages co-expressed CD163, which is consistent with the single-cell data (Fig. 2o, p). In addition, both CD68^+^ macrophages and MPO⁺ neutrophils were increased in the adventitia (data not shown). It is noteworthy that, in most forms of acute inflammation, monocyte and macrophage accumulation is generally observed 24 to 48 hours after onset ^27^. Indeed, in our mouse hindlimb ischemia model, macrophages were not yet increased 24 hours after induction of ischemia (data not shown). These findings also suggest that an event leading to increased macrophage accumulation in the media occurred prior to the onset of AD. We further characterized macrophages by multiplexed tissue imaging mass cytometry (IMC) and found that both CD163⁺MRC1(CD206)⁺ and CD163⁺MRC1(CD206)^neg∼low^, which are phenotypically consistent with AD-specific cytokine-expressing macrophages, accumulate in the media, with some located near CD31⁺ERG⁺ ECs (Fig. 2q and data not shown). Moreover, we confirmed that these CD163^+^ macrophages located near the pathological blood vessels do express VEGF and MMP9 protein (Fig. 2r). Taken together, these results suggest that cytokine-secreting CD163^+^MRC1(CD206)^neg∼low^ macrophages infiltrate into the media where they promote angiogenesis and modulate remodeling of ECM through VEGF and MMP9 before AD onset.

### AD Media are Hypoxic and Exhibit Microcalcification

Because hypoxia commonly drives angiogenesis ^28^, we investigated the hypoxic state of AD media by immunostainings for carbonic anhydrase-9 (CA9), a well-established marker of hypoxia ^29^. As hypothesized, we found high expression of CA9 in the SMCs and extracellular matrix around the middle part of the media (Fig. 3a, b). To investigate whether hypoxic changes in the media precede the onset of AD, we exposed human aortic SMCs (HAoSMCs) to hypoxic cell culture conditions. Consistent with the results of immunostainings, HAoSMCs began to express CA9 after only 24 hours under hypoxic condition, albite at low levels, which increased thereafter (Fig. 3c). These data support the notion that hypoxic responses in the media precede the onset of AD.

**Figure 3.**
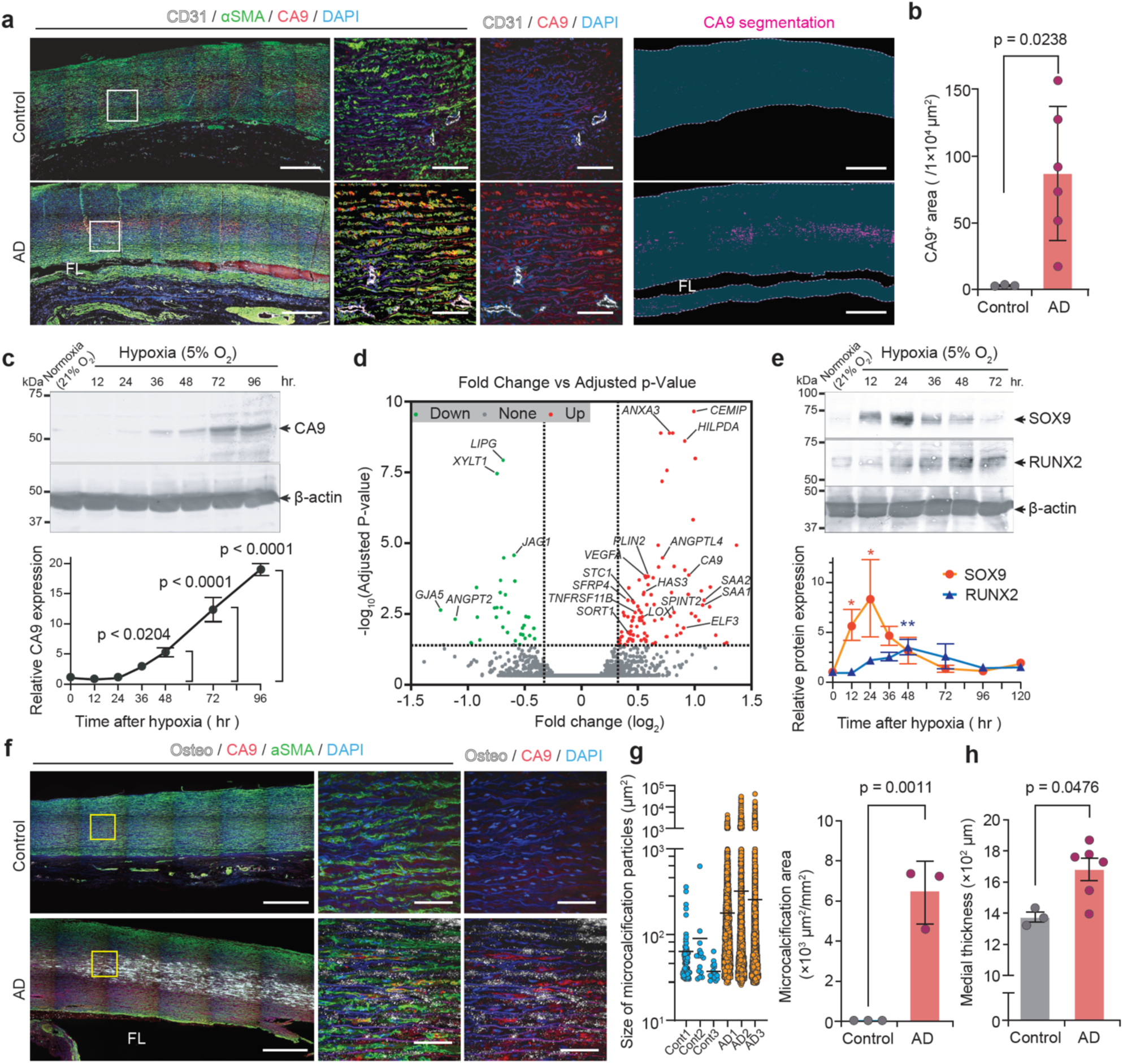
Hypoxia and Hydroxyapatite Deposition in the Aortic Media of Aortic Dissection. **(a)** Representative immunofluorescence images of aortic sections stained for CD31 (white), αSMA (green), and CA9 (red), counterstained with DAPI (blue). Magnified views of the regions indicated by the white boxes in the left panels are shown in the middle panels. Images of segmented CA9-positive area (purple) are shown in the right panels. FL, false lumen; the dotted line indicates the border of the media. Scale bars: 1 mm in wide-field images and 100 µm in magnified images. (**b**) Bar plot showing quantification of the CA9-positive area in the media. n = 3–6; data are expressed as mean ± S.D. (**c**) Representative western blot images showing CA9 and β-actin protein expression in human aortic smooth muscle cells (HAoSMCs) cultured under normoxic (21% O₂) or hypoxic (5% O₂) conditions for 96 hours. The graph shows densitometric analysis of CA9 protein levels. n = 3; data are expressed as mean ± SEM. (**d**) Volcano plot showing differentially expressed genes in HAoSMCs cultured under hypoxic conditions compared with normoxic conditions, based on bulk RNA-seq analysis. The dotted lines indicate a false discovery rate (FDR) of <0.1 and a fold change >1.25 or <0.80. Red dots indicate upregulated genes, and green dots indicate downregulated genes. (**e**) Representative western blot images showing SOX9, RUNX2, and β-actin protein expression in HAoSMCs cultured under normoxic (21% O₂) or hypoxic (5% O₂) conditions for 72 hours. The graph shows densitometric analysis of SOX9 and RUNX2 protein levels. n = 3; data are expressed as mean ± SEM. (**f**) Representative immunofluorescence images of aortic sections stained for OsteoSense (white), CA9 (red), and αSMA (green), counterstained with DAPI (blue). Magnified views of the yellow-boxed regions in the left panels are shown in the middle and right panels. FL, false lumen. Scale bars: 1 mm in wide-field images and 100 µm in magnified images. (**g**) Dot plots show the particle size of hydroxyapatite deposits in different sample groups (left). Bar plots show quantification of the area of hydroxyapatite deposits in the media (right). n = 3; data are expressed as mean ± S.D. (**H**) Bar plots show quantification of the thickness of media. n = 3–6; data are expressed as mean ± S.D.

We further investigated gene expression changes in HAoSMCs under hypoxia and identified 101 upregulated genes through bulk-RNAseq analysis (data not shown). Pathway analysis showed upregulation of several pathways including response to hypoxia (data not shown). Although the pathway analysis did not reveal a specific signature, we noticed increased expression of genes related to osteoblast differentiation, including *STC1*, *SFRP4*, and *ANGPTL4* (Fig. 3d and data not shown). We therefore explored the expression of transcription factors involved in osteochondrogenesis ^30^ and found that hypoxia induced the expression of RUNX2 (runt-related transcription factor 2) and SOX9 (Sry-related HMG box-9) at protein level (Fig. 3e and data not shown). More importantly, we indeed observed a significant deposition of hydroxyapatite in the media of AD samples, as assessed by the fluorescent bisphosphonate imaging probe OsteoSense (Fig. 3f, g, and data not shown), whereas calcium deposition, evaluated by Von Kossa staining, was only slightly higher or was similar in AD samples and controls (data not shown). Given that the aortic media in AD samples exhibited significant thickening (Fig. 3h), likely as a result of hypertension, our findings suggest that this structural change, at least in part, may contribute to the development of hypoxia, which in turn promotes tendency of SMCs to differentiate toward an osteochondrogenic phenotype.

However, since medial thickening due to hypertension is observed in many patients, it appears to be insufficient in itself to cause hypoxia, suggesting that additional factors are also involved.

### Blood vessels in the media are a risk factor for dissection

We next investigated whether these changes in the media could contribute to AD causation. To this end, we mapped the stiffness of media using atomic force microscopy (AFM). First, we evaluated the stiffness of areas in the middle to outer parts of the media (ROI-1), which contain CD31^+^ blood vessels, and in the luminal side (intima side; ROI-2) (Fig. 4a). Areas containing CD31^+^ blood vessels were significantly less stiff than inner parts in AD samples (Fig. 4b-d and data not shown). We further analyzed areas containing CD31⁺ blood vessels (ROI-2) and observed that the vessels and their perivascular regions exhibited greater softness compared to the surrounding tissue (Fig. 4e, f, and data not shown). Concentration of mechanical stress occurs when there are irregularities in the structural components; therefore, these results suggest that the increased blood vessels in the media form soft areas compared to the surroundings, and thus contribute to the development of AD.

**Figure 4.**
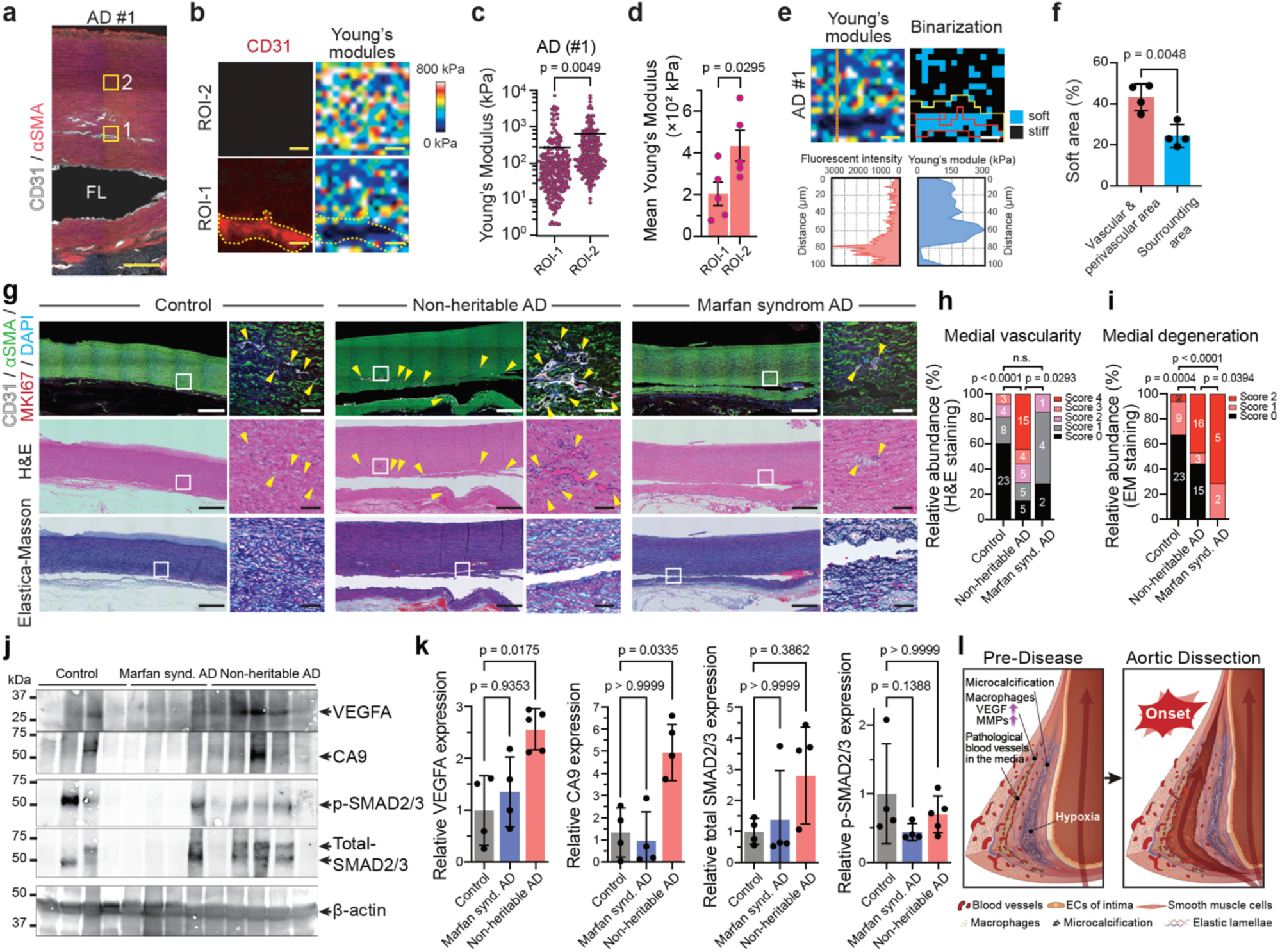
Stiffness of Pathological Medial Blood Vessels and Their Presence in Retrospective Specimens. **(a)** Representative image of stiffness measurement regions by atomic force microscopy (AFM). Region of interest 1 (ROI-1) includes CD31-positive blood vessels, while ROI-2 is located in the middle of the media. Sections were stained for CD31 (white) and αSMA (red). Scale bar: 500 μm. (**b**) Representative immunofluorescence images stained for CD31 (left panels), with corresponding Young’s modulus distribution maps for ROI-1 and ROI-2 shown in the right panels, as indicated in (**a**). The red dashed line indicates the margin of the CD31-positive blood vessel. Scale bar: 20 μm. (**c**) Dot plots show Young’s modulus measurements in the specified ROIs from one AD sample. (**d**) Bar plot showing quantification of the mean Young’s modulus in ROI-1 and ROI-2 across samples. n = 5; data are expressed as mean ± S.D. (**e**) The orange dashed line in the upper left panel indicates the region where fluorescence intensity and corresponding Young’s modulus were measured, as shown in the lower panels. The upper right panel shows a binary map of stiffness values divided by a specific threshold. The red line indicates the boundary of the CD31-positive blood vessel, and the yellow line indicates the boundary of perivascular area. (**f**) Bar plot showing the percentage of soft areas within the vascular and perivascular regions compared with the remaining area in ROI-1. n = 4; data are expressed as mean ± SEM. (**g**) Representative immunofluorescence and histological images of aortic sections from 38 non-dissected controls, 35 hyperacute aortic dissection cases (less than 12 hours post-onset), and 7 Marfan syndrome patients with dissection (retrospective specimens). Staining for CD31 (white), αSMA (green), Ki67 (red), and DAPI (blue) is shown in the upper panels. The middle and lower panels show hematoxylin and eosin (H&E) and Elastica-Masson (EM) staining, respectively. Magnified views of the white-boxed regions in the wide-field images are presented to the right. EM staining demonstrates medial degeneration. Arrowheads indicate blood vessels. Scale bars: 1 mm in wide-field images and 50 µm in magnified images. (**h**) Stacked bar graph showing the distribution of medial vascularity scores based on H&E staining across control, non-heritable hyperacute AD, and Marfan syndrome specimens. Each color represents a different score (0–4), and numbers indicate the number of samples in each category. (**i**) Stacked bar graph showing the distribution of medial degeneration scores based on EM staining across control, non-heritable hyperacute AD, and Marfan syndrome specimens. Each color represents a different score (0–2), and numbers indicate the number of samples in each category. (**j**) Representative western blot images showing VEGFA, CA9, phospho-SMAD2/3, total SMAD2/3, and β-actin protein expression in tissue lysates obtained from archived formalin-fixed, paraffin-embedded (FFPE) samples of control, non-heritable hyperacute AD, and Marfan syndrome specimens. n = 4–5. (**k**) Bar plot showing densitometric analysis of VEGFA, CA9, phospho-SMAD2/3, total SMAD2/3 protein levels. n = 4–5; data are expressed as mean ± SEM. (**l**) Schematic model illustrating the proposed mechanism of AD onset.

Lastly, we retrospectively evaluated an independent set of archived samples from 34 patients who underwent surgery for Stanford Type-A AD between 2018 and 2022. As expected, CD31 immunostaining of archived formalin-fixed paraffin-embedded (FFPE) specimens from AD patients showed significantly high vascularity than controls (Fig. 4g, and data not shown). Importantly, histopathological evaluation of 34 AD specimens by board-certified pathologists using hematoxylin and eosin (H&E) staining revealed a clear increase in medial blood vessels in 19 of these 34 AD cases (56%) (Fig. 4g, h, and data not shown). In contrast, AD samples obtained from patients with Marfan syndrome exhibited lower vascularity, as evaluated by CD31 immunostaining and H&E staining, as well as less migration of CD68^+^ macrophages, similar to that observed in controls (Fig. 4g, h, and data not shown).

When comparing non-heritable AD with AD associated with Marfan syndrome by immunostaining, a clear histological distinction was generally observed between the two. However, some Marfan syndrome samples exhibited mildly increased vascularity, while some non-heritable AD samples showed low vascularity, as observed by H&E staining. As a result, statistical analysis revealed only a modest overall difference (data not shown). Similarly, although the number of macrophages in the media differed markedly in most cases, one Marfan syndrome specimen also showed a modest increase in macrophages. Consequently, the difference in macrophage numbers did not reach statistical significance, likely due to the limited sample size (data not shown). On the other hand, Marfan syndrome samples exhibited significantly more elastic fiber fragmentation compared to both control and non-heritable AD specimens (Fig. 4g,i). We also confirmed higher expression of VEGF-A and CA9 proteins in non-heritable AD samples compared to those from AD associated with Marfan syndrome, while expression of phospho-Smad2/3, a critical intracellular pathway mediating TGF-β signaling commonly activated in heritable AD cases ^31^, was similar (Fig. 4j, k). Moreover, we found significantly higher vascularity in the adventitia of both non-heritable AD and Marfan syndrome samples compared to normal controls, with the former having the highest vascularity (data not shown). In addition, migration of inflammatory cells was not apparent in the media on H&E staining, but a significant increase was observed in the adventitia (data not shown). Taken together, these results suggest that angiogenesis and increased infiltration of macrophages into the media is observed in more than 50% of non-heritable AD and that the increased blood vessels in the media are likely involved in the onset of dissection (Fig. 4l).

## Discussion

In this study, we identified the angiogenic changes and a significant increase in blood vessels in the media that precede the onset of AD. Although it is not possible to obtain samples prior to the onset of AD, our time-course analysis revealed that hypoxia in the media, macrophage infiltration, hydroxyapatite deposition, and angiogenesis occur prior to dissection onset. Hypoxia induces VEGF-mediated angiogenesis and also triggers osteochondrogenic changes in SMCs. Hydroxyapatite promotes macrophage migration and alters their function ^32,33^. Macrophages are major actors in inducing angiogenesis and ECM remodeling, and are themselves capable of inducing osteogenic changes in vascular calcification ^34,35^. These processes may mutually reinforce each other, contributing to the formation of fragile, degenerated media with soft, pathological vessels.

While the precise cause of hypoxia remains unclear, contributing factors may include hypertension-induced medial thickening, atherosclerosis, and impaired blood flow or occlusion of the vasa vasorum ^36^. In the human thoracic aorta, oxygen diffusion from the lumen is estimated to reach only about 0.5 mm ^37^. When diffusion is insufficient, the region beyond this limit—especially the mid-media—is prone to hypoxia ^38^. Our findings support this, as the zone between the lumen and vasa vasorum supply showed hypoxia, hydroxyapatite deposition, and neovessel extension into this region. Although the timing of these changes remains unclear, this structurally compromised zone may be particularly vulnerable to stress, such as sudden blood pressure changes. This vascular increase cannot be reproduced in standard mouse models, likely due to differences in aortic thickness and oxygen diffusion ^39^. Therefore, at present, this phenomenon can only be reliably examined with human samples. To determine when these changes begin to occur, it is necessary to develop a model that replicates the human condition. The adventitia and perivascular adipose tissue are also involved in vascular inflammation and remodeling ^40^, so the increase in blood vessels in these areas may additionally contribute to the onset. Regarding calcification, we speculate that hypoxia, together with imbalances in calcium and phosphate, contributes to medial microcalcification. Macrophages, in addition to SMCs, may play a significant role, particularly as potent inducers of osteogenic transdifferentiation in SMCs ^41^. Further investigation into these mechanisms is warranted.

AA is a well-known chronic disease characterized by degeneration and remodeling of the aortic wall. The pathological changes in aneurysmal walls have been extensively studied, and it is well established that the disease process involves infiltration and activation of immune cells such as macrophages, activation of fibroblasts, phenotypic modulation of smooth muscle cells, and angiogenesis. In contrast, while cystic medial necrosis is a known histological feature in heritable connective tissue disorders associated with dissection, such as Marfan syndrome, non-heritable acute AD is generally considered to arise from histologically normal aortic walls. Therefore, the concept of “medial degeneration” as a prerequisite or predisposing condition for AD remains controversial ^42,43^. Our findings suggest that key cellular players involved in aneurysm formation may also contribute to the initiation of AD. Although AD and AA were historically conflated—an issue further compounded by murine models in which aneurysm and dissection frequently develop simultaneously—they are now recognized as distinct clinical and pathological entities. In particular, acute AD typically arises independently of preexisting aneurysms and follows a fundamentally different disease trajectory. Nevertheless, our data imply that both conditions may share common early cellular responses within the aortic wall. It is therefore conceivable that additional factors determine whether these initial changes lead to chronic aneurysm formation or to an acute dissection event. Identifying such a branching point would have important implications for both mechanistic understanding and clinical intervention.

Our findings also suggest that non-heritable AD, which accounts for the majority of cases, can be classified into angiogenic and non-angiogenic subtypes. While many cases showed increased medial vasa vasorum, others did not. Interestingly, some angiogenic cases lacked tip and proliferative ECs by immunostaining and did not show clear angiogenic signatures in scRNA-seq, although immature EC clusters were present. These may represent a quiescent subtype, and the distinction may reflect angiogenic maturation or timing. Determining the proportion of each subtype and their relationship to macrophage infiltration and calcification will be important in future research. Moreover, given our limited sample size, we could not distinguish patient-profile differences between subtypes; assessing variations in past medical history and lifestyle factors (e.g., diet and physical activity) remains an important goal. Significantly, increased vascularity can be detected using H&E staining, a widely available clinical method, suggesting the feasibility of broader diagnostic application. Whether these angiogenic subtypes differ in clinical outcomes remains to be investigated. Few studies have investigated the relationship between angiogenesis and acute AD. A research group from Finland reported aortic wall remodeling accompanied by increased vascularization and chronic inflammation in the ascending aorta ^44^. In addition, one report from Italy described increased vasa vasorum ^45^. Angiogenesis in AD may therefore represent a phenomenon common across populations. Comparisons between non-heritable AD and Marfan syndrome with acute revealed distinct histological features, with angiogenesis largely absent in Marfan cases. This strongly suggests different molecular mechanisms underlie each form. Taken together, our results support a new model in which more than 50% of non-heritable AD arises as an angiogenic disease of the aortic media. Although prediction remains difficult, detection of medial angiogenesis may serve as a new avenue for risk assessment. Targeting this angiogenesis and relieving hypoxia may offer future strategies for prevention and treatment.

## Acknowledgements

We thank Ms. Y. Mitani, Ms. C. Hirose, Ms. T. Yuhi and Ms. M. Oura for their technical assistance. We also appreciate the technical support provided by the Division of Advanced Biomedicine, Medical Research Support Center, Graduate School of Medicine, Kyoto University, for the IMC study, the NGS core facility at the Research Institute for Microbial Diseases of Osaka University for the bulk RNA sequencing and data analysis, and Biken Biomics for the single cell RNA sequencing. We acknowledge the following financial support for the research presented in this article: the Japan Agency for Medical Research and Development (AMED) (AMED-PRIME JP21gm6210009, AMED-FORCE JP24gm4010023), the KAKENHI (24K02221, 24K10024, 22H05063, 22H05060, 20H03435), the Mitani Foundation for Research and Development, the Takeda Science Foundation, the Kato Memorial Fund for Incurable Disease Research, and the Kanazawa University SAKIGAKE project.

## Author contributions

H.N. conceived the study. H.N and K.Y. designed the experiments. K.Y. performed immunostaining, cell assays, protein assays, RNAseq analysis and analyzed most of the data with the help of T.M, A.M. and B.H. K.I., Y.Y. and H.T. performed surgery and collected samples. Y.I., A.G. and D.M. performed pathological evaluation. T.I performed cell isolation. H.N and T.I. performed single cell analysis. T.I. analyzed the single cell analysis data. T.M. N.S. H.N. and H.Y. performed AFM. H.N. M.H. and T.W. performed IMC. T.S. helped with single-cell data interpretation. K.Y. and H.N. wrote the manuscript. H.N. edited the final version of the manuscript, with input from all authors. H.N acquired the funding. H.A. and H.T. provided critical inputs to the manuscript. H.N supervised all the research. All the authors read and approved the final manuscript.

## Competing interests

The authors declare no competing interests.

**Correspondence and requests for materials** should be addressed to Hisamichi Naito as the primary contact.

## Materials and Methods

### Ethics statements

The collection of human samples and research conducted in this study were approved by the Ethics Committee of Kanazawa University (approval number: 2021-245 (113884)). Written informed consent for collection and research use of the surgically removed aorta was obtained from each patient prior to the operation. All the animal experimental protocols were approved by the Animal Care and Use Committee of Kanazawa University.

### Acquisition and processing of human aorta, and cell isolation

For AD patients, ascending aorta (zone 0) was obtained by total aortic arch replacement. Immediately after resection, samples were washed with saline and stored in MACS Tissue Storage Solution (Miltenyi Biotec, Bergisch Gladbach, Germany) at 4°C. For the immunostainings, the aorta was fixed in 10% formalin or 4% paraformaldehyde (PFA). For cell sorting, the outer layers (comprising the outer part of the dissected media and adventitia) were separated from the inner layers (comprising the inner part of the dissected media and intima), and tissue pieces ranging from 10 to 25 cm² were minced into small fine pieces (1 mm or smaller). The tissue was then digested, and a single-cell suspension was prepared as previously described with some modifications ^46^. Briefly, an enzyme cocktail was prepared containing 3 mg/ml collagenase type II (LS004176, Worthington), 0.15 mg/ml collagenase type XI (H3506, Sigma), 0.25 mg/ml soybean trypsin inhibitor (LS003571, Worthington), 0.1875 mg/ml lyophilized elastase (LS002292, Worthington), and 0.24 mg/ml hyaluronidase type I (H4034, Sigma) in 4% FBS/PBS supplemented with 1 mM CaCl_2_. Digestion was carried out at 37°C for 60 minutes with gentle agitation (30 rpm). The resulting tissue digest was passed through 100-µm and 40-µm cell strainers (BD Corning), centrifuged at 330 x g for 5 minutes.

Erythrocytes were lysed with ACK Lysing Buffer (Lonza Japan, Tokyo, Japan) and resuspended in 4% BSA/PBS. Cells were first treated with Fc block (BioLegend #422302) to inhibit non-antigen-specific bindings and then cell-surface antigen staining was performed for CD31 (clone WM59, BioLegend) and CD45 (clone HI30, BioLegend). The stained cells were analyzed, and CD31^+^CD45^-^ cells were sorted using a FACS Aria Fusion (BD Biosciences) or CytoFLEX SRT (Beckman Coulter, Brea, CA, USA).

### Single-Cell RNA Sequencing

Single-cell capturing and downstream library construction were performed using the Chromium Next GEM Single Cell 3’ Kit v3.1 for Dual Index (PN-1000269, 10x Genomics) according to the manufacturer’s protocol. Briefly, single-cell suspensions obtained from fresh patient samples were diluted to 1000 cells/μl in 0.1% BSA/PBS. A single-cell suspension, barcoded gel beads, and partitioning oil were loaded into Chromium Chip G (PN-1000127, 10x Genomics) to generate Gel Beads in Emulsion (GEMs). The transcripts were reverse transcribed within the GEMs, and the resulting cDNA was amplified with the barcodes. Library construction involved fragmentation, adapter ligation, and a sample index PCR. The constructed libraries were sequenced using the DNBSEQ-G400 (MGI Tech, Shenzhen, Guangdong, China) at Biken Biomics Co., Ltd. (Osaka, Japan), with 28 bp x 100 bp paired-end, dual indexing. For demultiplexing and mapping of the sequenced data to the GRCh38 reference genome (ensemble version 98), CellRanger software version 6.1.2 (10x Genomics) was used.

### Single-cell RNA-seq Data Processing and Integration

Single-cell RNA-seq data were processed and analyzed using the Seurat package (version 4.4.0). The percentage of mitochondrial UMI counts was calculated using the Seurat function *PercentageFeatureSet*. Cells with >15% mitochondrial UMI or >8,000 detected features were excluded from further analysis. Gene expression matrices were normalized using the *LogNormalize* function with default parameters. Cell cycle regression was performed using the *CellCycleScoring* function, which assigns S and G2/M scores and classifies cells into G1, S, or G2/M phases. For data integration and batch correction, two complementary workflows were used. In the first, highly variable genes (top 2000) were identified with *FindVariableFeatures*, and unwanted variation - including mitochondrial UMI counts, number of RNA features, total RNA counts, and cell cycle scores - was regressed out using *ScaleData*.

Principal component analysis (PCA) was conducted with *RunPCA*, followed by integration using *RunHarmony*. The optimal number of Harmony dimensions was determined using a modified ElbowPlot method (Harvard Chan Bioinformatics Core). Dimensionality reduction and clustering were performed using *RunUMAP*, *FindNeighbors*, and *FindClusters*. In the second workflow, data were normalized and technical and biological variation was regressed using *SCTransform*. Integration features were selected with *SelectIntegrationFeatures*, and PCA and subsequent steps were performed as in the first workflow. Cell-cell interaction analyses were conducted using CellPhoneDB (version 5.0.1) and CellChat (version 1.6.1), with visualization of CellPhoneDB results carried out using the ktplots package. The complete analysis pipeline for single-cell RNA-seq data processing and integration, including all code and parameters, <u>will be available on GitHub.</u>

### Immunostaining of aorta

Formalin-fixed paraffin-embedded (FFPE) tissue were sectioned in 4 µm slices, using a sliding microtome (Retoratome, Yamato Kohki Industrial, Saitama Japan), mounted onto microscope slides, and processed for staining with the antibodies and reagents (details available upon request). Immunostaining was performed as previously described ^47^. Briefly, the sections were washed sequentially in xylene, ethanol, and PBS; heat-induced epitope retrieval was then performed in a pressure cooker SP-5D152 (Siroca, Tokyo, Japan) for 5 min in DAKO Target Retrieval Solution (pH 9.0, #S2367 or pH 6.1, #S1699, Agilent) followed by cooling to room temperature prior to staining. Cryosections of 4% PFA-fixed OCT (Sakura Finetek, Tokyo, Japan)-embedded tissue was cut at 20 µm, mounted on MAS-coated slide glass (Matsunami Glass, #S9115), and washed in PBS. These sections were blocked with DAKO Protein Block (Agilent, #X0909) and incubated overnight at 4°C with a mixture of primary antibodies diluted in the SignalStain Antibody Diluent (CST, #8112). Primary antibodies used for immunofluorescence analysis were anti-CD31 (Agilent DAKO or rabbit IgG, CST #77699), anti-α smooth muscle actin (αSMA) (Biolegend #904601), anti-CD68 (Agilent DAKO #M0876), anti-CD163 (CST #93498), anti-APLNR (Proteintech #25800-1-AP), anti-ESM1 (Abcam #ab224591), anti-Ki67 (Agilent DAKO #, MIB-1) or anti-Ki67 (CST #9027), anti-VEGFA (ABclonal #A0280), anti-MMP9 (CST #13667), anti-CA9 (Abcam #ab270401); primary antibody concentrations were optimized through testing of various dilutions to achieve optimal staining with minimal background, and negative controls were performed by omitting the primary antibodies. The following secondary antibodies were used: goat anti-mouse IgG1-Alexa 647 (A-21240), goat anti-rabbit IgG-Alexa 647 (A-21244), goat anti-mouse IgG1-Alexa 568 (A-21124), goat anti-mouse IgG2a-Alexa 568 (A-21134), goat anti-mouse IgG2b-Alexa 568 (A-21148), goat anti-rabbit IgG-Alexa 568 (A-11011), goat anti-rat IgG-Alexa 568 (A-11077), goat anti-mouse IgG1-Alexa 488 (A-21121), goat anti-mouse IgG2a-Alexa 488 (A-21134), goat anti-mouse IgG3-Alexa 488 (A-21151), and goat anti-rabbit IgG-Alexa 488 (A-11034), all from Thermo Fisher Scientific (Molecular Probes, Inc.). Following secondary antibody incubation, sections were incubated for 15 minutes at room temperature in DAPI solution (Thermo Fisher Scientific, Molecular Probes, Inc., #D1306) and then mounted with FluoroMount (Diagnostic BioSystems, #K024). Microcalcification detection was performed using the IVISense Osteo 680 Fluorescent Probe (OsteoSense, Revvity, #NEV10020EX). Laser scanning confocal microscopy was carried out by acquiring tile scans on an Andor Dragonfly 200 system (Oxford Instruments, Abingdon-on-Thames, UK) equipped with Fusion Stitcher software and built on a Nikon ECLIPSE Ti2 inverted microscope (Nikon Corporation, Tokyo, Japan). Images were acquired under identical microscope settings using sequential acquisition of different channels to prevent signal overlap between Alexa fluorophores, and non-overlapping fluorophores were selected for colocalization analysis. All images were saved as non-compressed TIFF files and quantified using AIVIA software (Leica Microsystems, Wetzlar, Germany). For quantification, CD31^+^ ECs within the medial area were counted across the entire section and expressed as the number of CD31^+^ ECs per actual medial area.

### Imaging mass cytometry

Imaging mass cytometry of FFPE sections was performed for macrophages and EC profiling. FFPE tissues were sectioned at 4 µm thickness, mounted onto microscope slides, and processed for staining with the antibodies (details available upon request). In brief, sections were deparaffinized and stained with anti-CD163-147Sm (#3147021D, Standard BioTools, South San Francisco, CA, USA), anti-collagen type I-169Tm (#3169023D, Standard BioTools), anti-Fibronectin-174Yb (#91H034174, Standard BioTools), anti-Alpha-Smooth Muscle Actin-209Bi (#3209017D, Standard BioTools), anti-CD206 (#87887SF, Cell Signaling Technology, Danvers, MA, USA) labeled with 141Pr (#201141A, Standard BioTools), anti-CD31 (#AF3628, R&D Systems) labeled with151Eu (#201151A, Standard BioTools), anti-ERG (ab214796, abcam) labeled with 153Eu (#201153A, Standard BioTools), anti-CD86 (#76755SF, Cell Signaling Technology) labeled with 158Gd (#201158A, Standard BioTools), anti-CD68 (#26042SF, Cell Signaling Technology) labeled with 159Tb (#201159A, Standard BioTools), anti-Ki67 (#87887SF, Cell Signaling Technology) labeled with 165Ho (#201165A, Standard BioTools), anti-CD14 (#43878SF, Cell Signaling Technology) labeled with 168Er (#201168A, Standard BioTools) and anti-Myeloperoxidase (#88757, Cell Signaling Technology) labeled with 175Lu (#201175A, Standard BioTools), following the manufacturer’s protocol. The tissue slide was ablated using a Hyperion Imaging System (Standard BioTools). AI-powered cell segmentation was performed using AIVIA software (Leica Microsystems).

### Western blotting

For in vitro cultured cells, human aortic smooth muscle cells (HAoSMCs, Lonza #CC-2571) were mechanically homogenized in Pierce RIPA Buffer (Thermo Scientific, #89900) supplemented with a protease inhibitor cocktail (Complete Mini, Roche, #04693159001). Human aortic tissue FFPE specimens were homogenized using the Qproteome FFPE Tissue Kit (Qiagen, #37623). Lysates were centrifuged at 20,000 x g for 15 minutes at 4°C, and the supernatant was used for subsequent experiments. Protein samples were separated by SDS-PAGE on 8-15% Bis-Tris acrylamide gels under reducing conditions (Mini-PROTEAN II system, Bio-Rad) and transferred to a polyvinylidene difluoride (PVDF) membrane (Millipore, #IPVH00010) using the Trans-Blot Turbo Transfer System (Bio-Rad). The membrane was blocked for 30 minutes at room temperature with PVDF Blocking Reagent (TOYOBO, #NYPBR01). Primary antibodies were then incubated overnight at 4°C in antibody diluent buffer (Can Get Signal, TOYOBO, #NKB-101), followed by incubation with alkaline phosphatase (AP)-conjugated secondary antibodies (CST, #7056 for anti-mouse IgG and #7057 for anti-rabbit IgG) for 1 hour at room temperature in the same diluent. The primary antibodies used were as follows: anti-CA9 (Abcam, #ab270401), anti-SOX9 (CST, #82630), anti-RUNX2 (CST, #12556), and anti-VEGFA (ABclonal, #A0280). Blotting for β-actin (WAKO Fujifilm, #010-27841) or GAPDH (WAKO Fujifilm, #010-25841) was performed as a loading control. The membrane was developed using the BCIP/NBT substrate kit (Sigma, #B5655) for an appropriate duration, and band densities were quantified using LI-CORbio Image Studio software (LICORbio, Nebraska, USA).

### Mouse hind-limb ischemia model

The ischemic hind-limb model was established using ten-week-old male C57BL6/J mice (SLC, Japan). Hindlimb ischemia was induced as previously described ^48^. Briefly, the left femoral artery and vein were exposed and resected from the proximal portion near the inguinal ligament to the distal portion of the saphenous artery, with the remaining arterial branches excised. The contralateral hind-limb served as an internal control. Blood flow in the left (ischemic) and right (non-ischemic, contralateral) hind-limbs was measured by laser Doppler imaging (Omegawave, Japan), and the average of 10 measurements was used to calculate the ratio of blood flow between the ischemic and contralateral sides at each time point. The gastrocnemius muscles were dissected and fixed in 4% PFA. Fixed samples were either embedded in OCT compound and sectioned at 20 µm for histological analysis or used for 3D immunostaining. For the assessment of capillary density, both non-ischemic and ischemic muscle sections were stained with an anti-CD31 (Biolegend, #102502) antibody. Immunostained images were acquired using confocal microscopy, and histological assessments were performed on five randomly selected fields per tissue section. For the 3D immunostaining, tissues were prepared and stained as previously described ^49^. Briefly, tissues were first delipidated and decolorized using CUBIC-L (T3740, Tokyo Chemical Industry, Tokyo, Japan).

Staining was performed over 7 days with primary IgG antibodies and Alexa Fluor-labeled Fab fragments (Jackson ImmunoResearch, Pennsylvania, USA). After washing, the tissues were refractive index (RI)-matched using CUBIC-R+(M) (T3741, Tokyo Chemical Industry) according to the manufacturer’s instructions. Images were acquired using a Lightsheet 7 microscope (Carl Zeiss, Oberkochen, Germany) and analyzed with Arivis Pro software (Carl Zeiss).

### Pathological Evaluation of Specimens

A total of 34 specimens of Stanford type A non-hereditary hyperacute AD and 7 specimens from patients who developed acute aortic dissection and were subsequently diagnosed with Marfan syndrome, all surgically treated at Kanazawa University, were analyzed using H&E and EM staining. Thirty-eight specimens from autopsies of patients with other diseases at Akita University were used as controls. The evaluations were performed by two board-certified pathologists. Vessel size in the aortic media was graded as “Vasodilatation” on a three-tier grading 0 to 2, with 2 indicating the largest vessels, and the number of vessels was assessed as “Vascular count” on a three-point scale, with 2 representing the highest vessel count. The extent of inflammatory cell infiltration in the media was graded as “Inflammation” on a three-point scale, with 2 indicating the greatest degree of inflammatory cell infiltration, and the condition of the elastic fibers was assessed as “Degeneration” on a three-point scale, with 2 indicating the most severe degeneration. The sum of the Vasodilatation and Vascular count scores was defined as the Vascularity score (not shown).

### Cell culture and hypoxic treatment

HAoSMCs were maintained in SmGM-2 Smooth Muscle Cell Growth Medium-2 BulletKit (Lonza #CC-3182) at 37°C in a humidified atmosphere containing 5% CO_2_. Cells were grown to semi-confluence and used at passages 4 to 6.

Experiments were performed using two cell lines derived from different donors. For hypoxia experiments, 5% O_2_ hypoxic conditions were achieved by placing the cells in a Multigas incubator (MCO-50M, PHCbi, Japan) connected to two gas cylinders containing CO_2_ and N_2_.

### Bulk RNA-seq analysis

RNA sequencing was performed on HAoSMCs that were exposed to 5% O₂ for 48 hours prior to RNA isolation. Total RNA was extracted using the RNeasy Mini Kit (Qiagen, Hilden, Germany), and RNA-seq libraries were prepared using the TruSeq Stranded mRNA Library Prep Kit (Illumina) according to the manufacturer’s protocols. Sequencing was conducted on a NovaSeq X Plus sequencer (Illumina) in paired-end mode (2 × 101 nt). After adapter trimming with Trimmomatic, the reads were aligned to the hg38 reference genome using HISAT2 version 2.1.0. Gene expression levels were quantified as fragments per kilobase of exons per million mapped fragments (FPKMs) using Cufflinks version 2.2.1. Data were analyzed with iDEP 2.01. FDR cutoff was set at 0.1 and the minimum fold-change was set at 1.25. Pathway analysis was performed using clusterProfiler package ^50^.

### Quantitative reverse transcription PCR

RNA was extracted using RNeasy Mini Kits (Qiagen), and cDNA generated using reverse transcriptase from the PrimeScript RT reagent Kit (Takara, Otsu, Japan). Real-time PCR was performed using a Thermal Cycler Dice Real Time System III (Takara). The polymerase chain reaction was performed on cDNA using specific primers. The sequences of the gene-specific primers were as follows:

GAPDH 5’-TGG CAA AGT GGA GAT TGT TGC C-3’

5’-AAG ATG GTG ATG GGC TTC CCG-3’

SOX9 5’-GAG CAG ACG CAC ATC TC-3’

5’-CCT GGG GAT TGC CCC GA-3’

RUNX2 5’-CCC GTG GCC TTC AAG GT-3’

5’-CGT TAC CCG CCA TGA CAG TA-3’

STC1 5’-TGA GGC GGA GCA GAA TGA CT-3’

5’-CAG GTG GAG TTT TCC AGG CAT-3’

SAA1 5’-AAC TAT GAT GCT GCC AAA AGG-3’

5’-TGG ATA TTC TCT CTG GCA TCG-3’

ELF1 5’-CCT TAC AGT TGA AGC TTC TTG TCA T-3’

5’-GGG GCA ACA ACC ATG TCA TC-3’

SFRP4 5’-TCT ATG ACC GTG GCG TGT GC-3’

5’-ACC GAT CGG GGC TTA GGC GTT TAC-3’

ANGPTL4 5’-GCA GGA TCC AGC AAC TCT TC-3’

5’-GGT CTA GGT GCT TGT GGT CC-3’

### Mechanical Characterization of the Aorta by Atomic Force Microscopy

Mechanical characterization of the aorta was performed using atomic force microscopy (AFM; NanoWizard 3, JPK Instruments, Berlin, Germany) in combination with confocal fluorescence microscopy (C2, Nikon Instruments, Tokyo, Japan). Fresh aortic tissue was embedded in OCT compound and snap-frozen using an isopentane/dry ice mixture. The samples were sectioned at 40 µm thickness and mounted on MAS-coated slides (Matsunami Glass, #S9115). Sections were then fixed with 4% PFA in phosphate buffer (PB) for 5 minutes at room temperature, followed by immunostaining for CD31 and αSMA as previously described. Silicon cantilevers with a spring constant of *k* ∼ 0.3 N/m and a conical tip (CSC37, μmasch Inc., Tallinn, Estonia) were used. Force-indentation (*f–i*) curves were acquired from sections in PBS solution. The spatial distribution of Young’s modulus (*E*) was determined according to previously described protocols ^51^. In brief, *E* was calculated from the approach curves using the Hertzian equation:

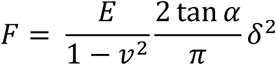

where *F* is the loading force (35 nN), *δ* is the indentation depth, *ν* is the Poisson’s ratio, and α is the semi-vertical angle of the indenter ^52^. In this study, *ν* and α were assumed to be 0.5 and 20°, respectively. Young’s modulus (*E*) was determined by fitting the data in the indentation depth range of 1–5 μm using the JPK data processing software (version 6.4.27). Binary stiffness maps were generated by dividing each sample into soft and stiff regions using a threshold defined as the lower one-third of the stiffness values measured within that sample. The vascular area was defined based on CD31 staining, and the perivascular area was designated as a 2-pixel-wide region surrounding the vascular area(1 pixel = 6.25 μm).

## Statistical analysis

All statistical analyses were performed using Prism 10 (GraphPad Software). All data are presented as the mean ± standard deviation (S.D.) or standard error of the mean (SEM). Differences between values were examined using two-tailed Mann-Whitney U tests. For comparisons among two or more groups, one-way ANOVA with a Tukey post-hoc test was used for parametric variables, the Kruskal-Wallis test for non-parametric variables, and the Fisher’s exact test for categorical variables. A two-sided p-value of less than 0.05 was considered statistically significant.

## Data and materials availability

The data supporting the findings of this study are available within the article and its Supplementary Information. The scRNA-seq data of human aorta and the RNA-seq data of HAoSMCs generated in this report will be deposited in GEO under accession number **GSE297281** and **GSE298721**.

## Competing interests

The authors declare no competing interests.

## Notes

### Competing Interest Statement

The authors have declared no competing interest.

